## Supplemental Figures for "Aβ-driven nuclear pore complex dysfunction alters activation of necroptosis proteins in a mouse model of Alzheimer’s Disease"

### SUPPLEMENTARY FIGURES

#### S1 *App* KI and WT neurons have comparable levels of pTau

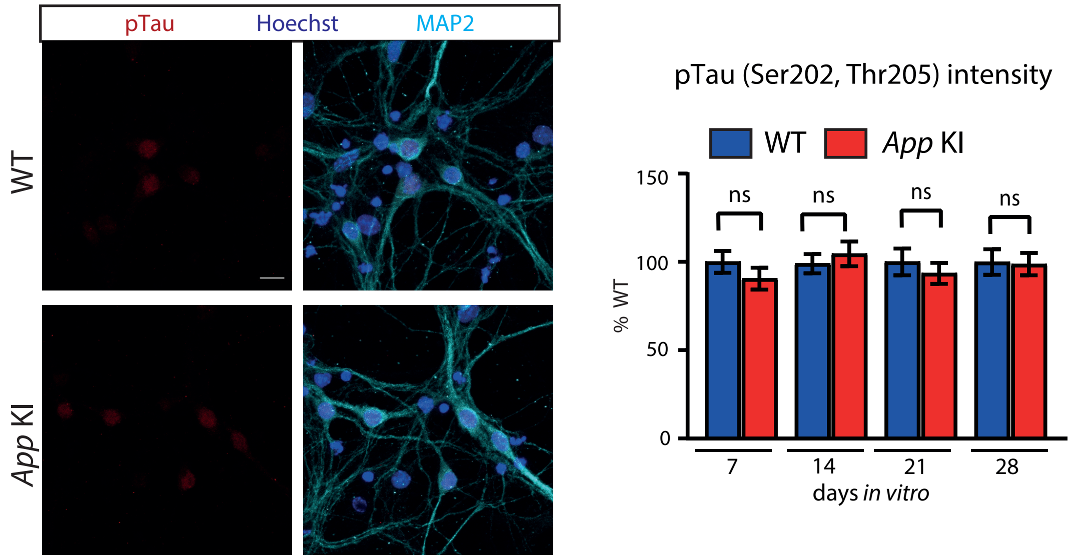


#### S2 *App* KI and WT cocultures show low levels of apoptosis and necrosis at basal state

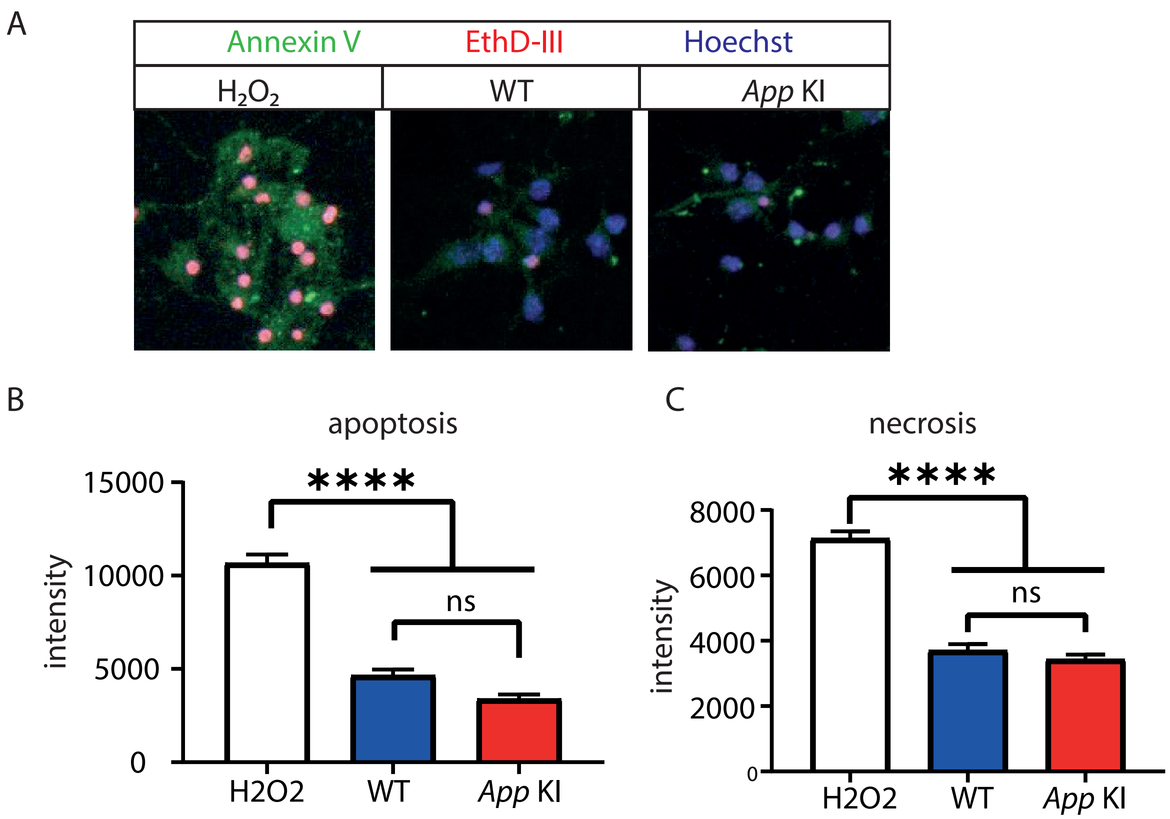

### SUPPLEMENTARY FIGURES

#### S3 RL1 positive fluorescent puncta on the nucleus colocalizes with NUP98

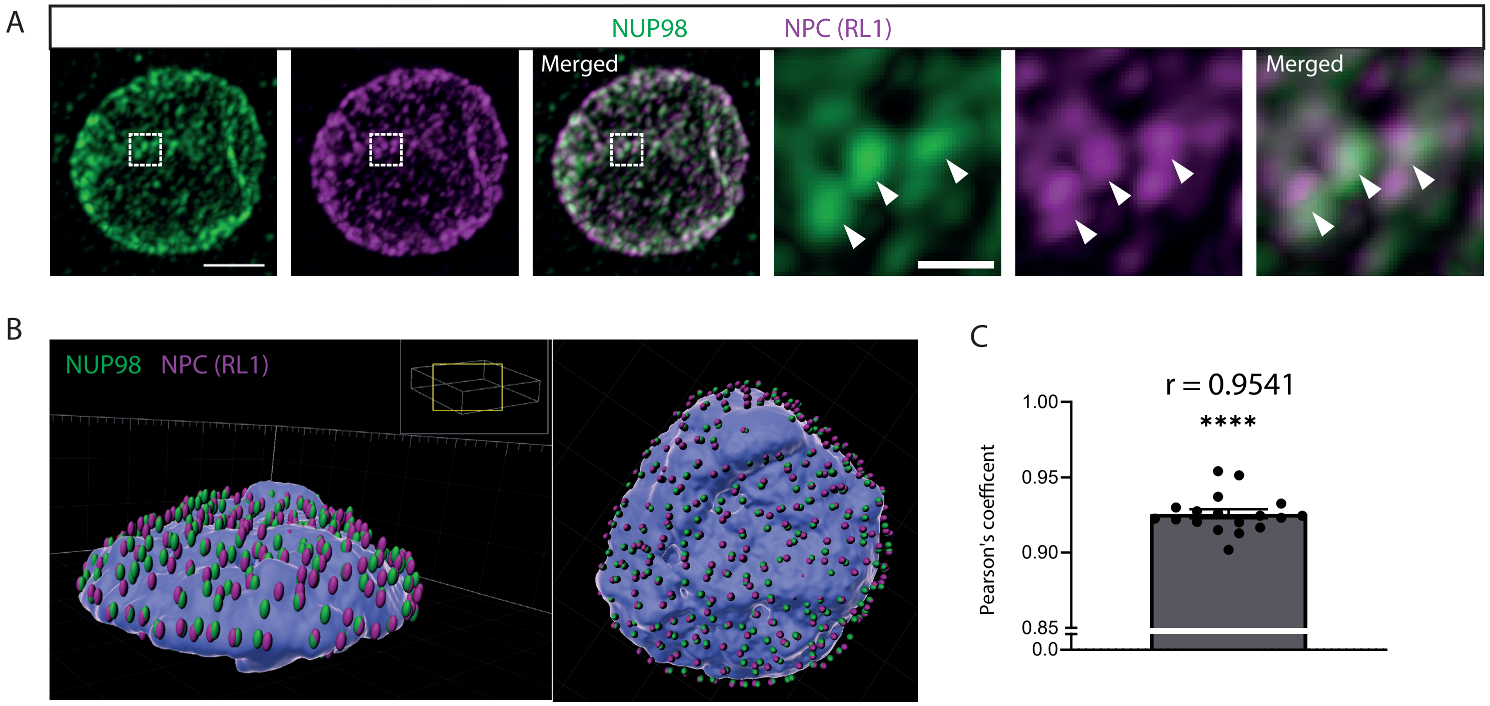


#### S4 The sphericity index and LAMIN-B1 expression in the nucleus is comparable in *App* KI and WT neurons

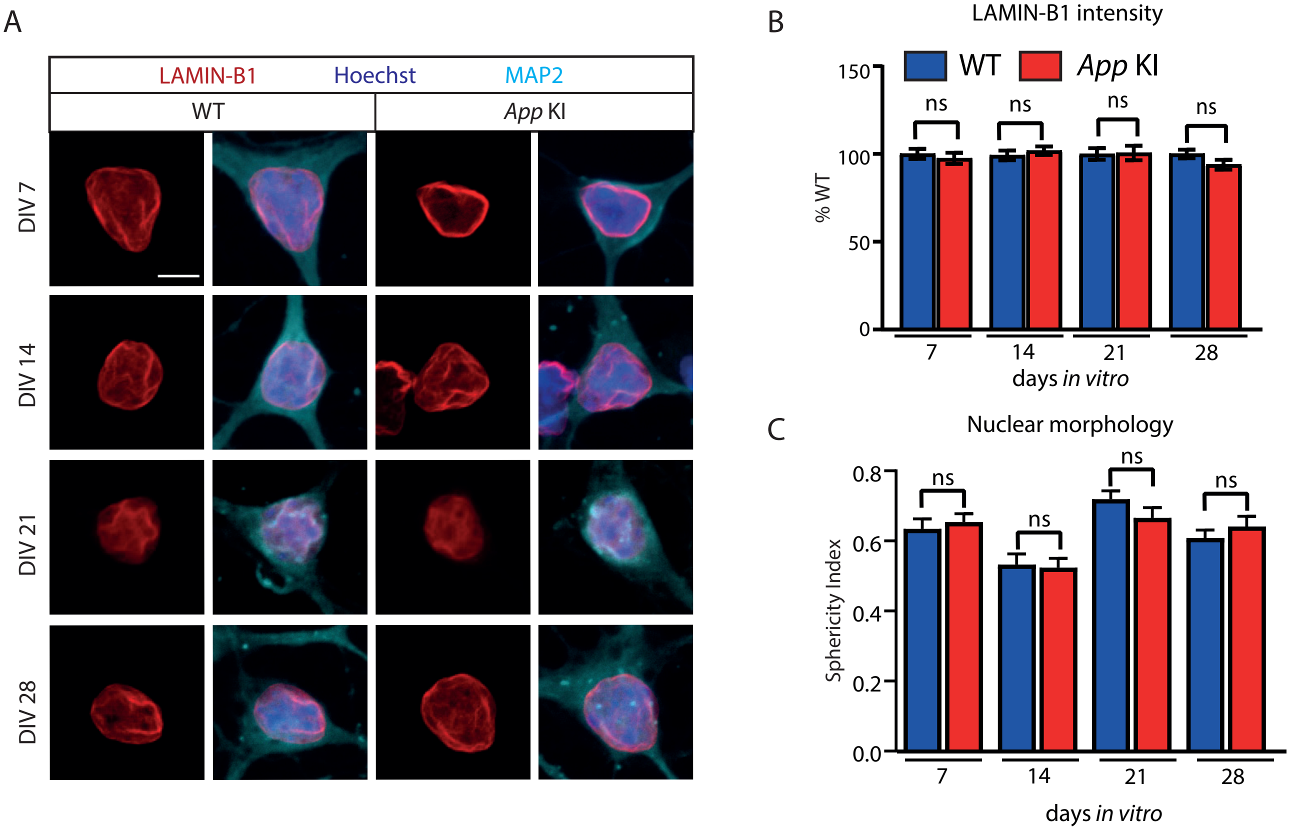

### SUPPLEMENTARY FIGURES

#### S5 Labeling of *App* KI neuronal nuclei with pan-NPC mAB414 antibodies reveal similar deficits

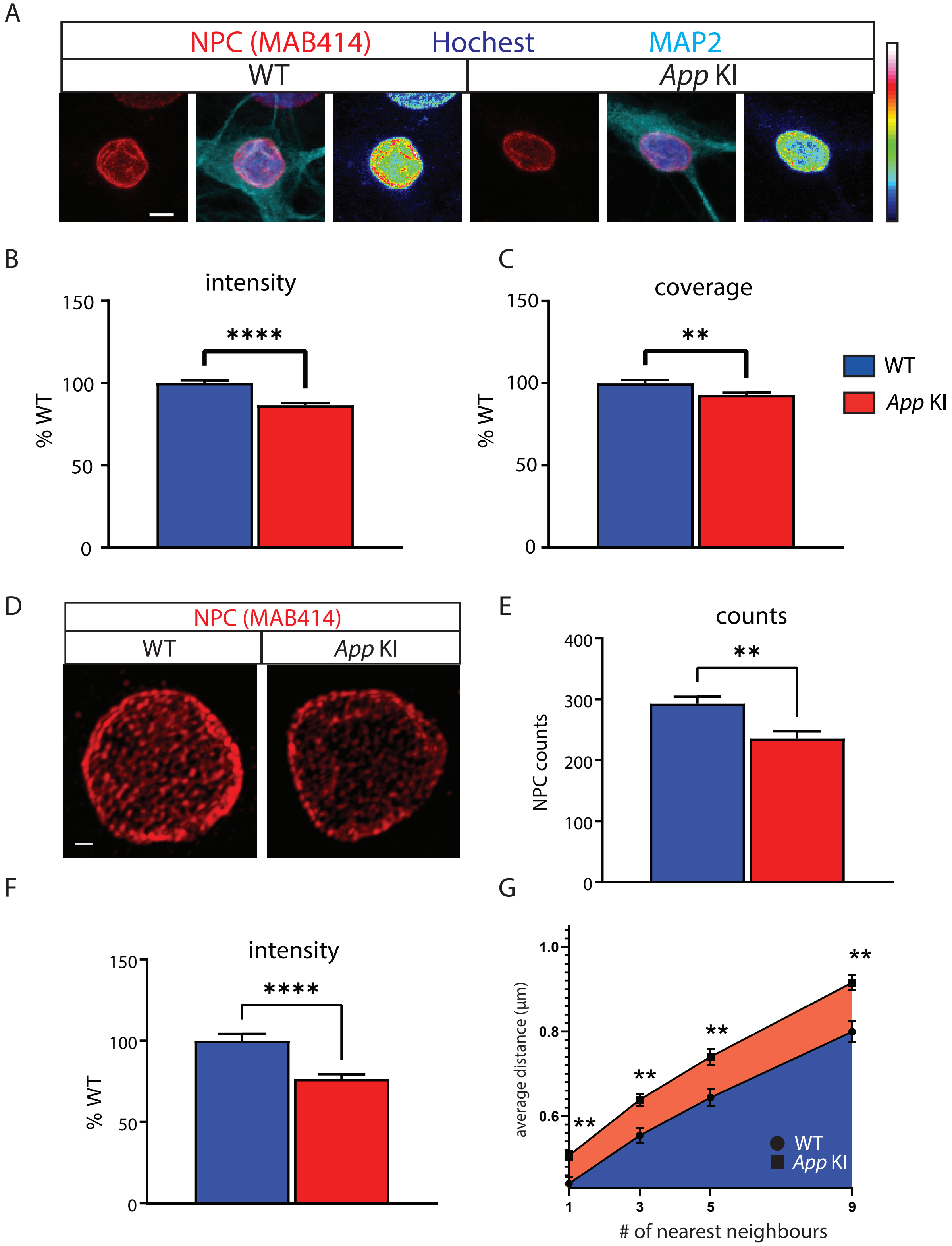

### SUPPLEMENTARY FIGURES

#### S6 Reduction of NUP98 positive puncta in *App* KI nuclei

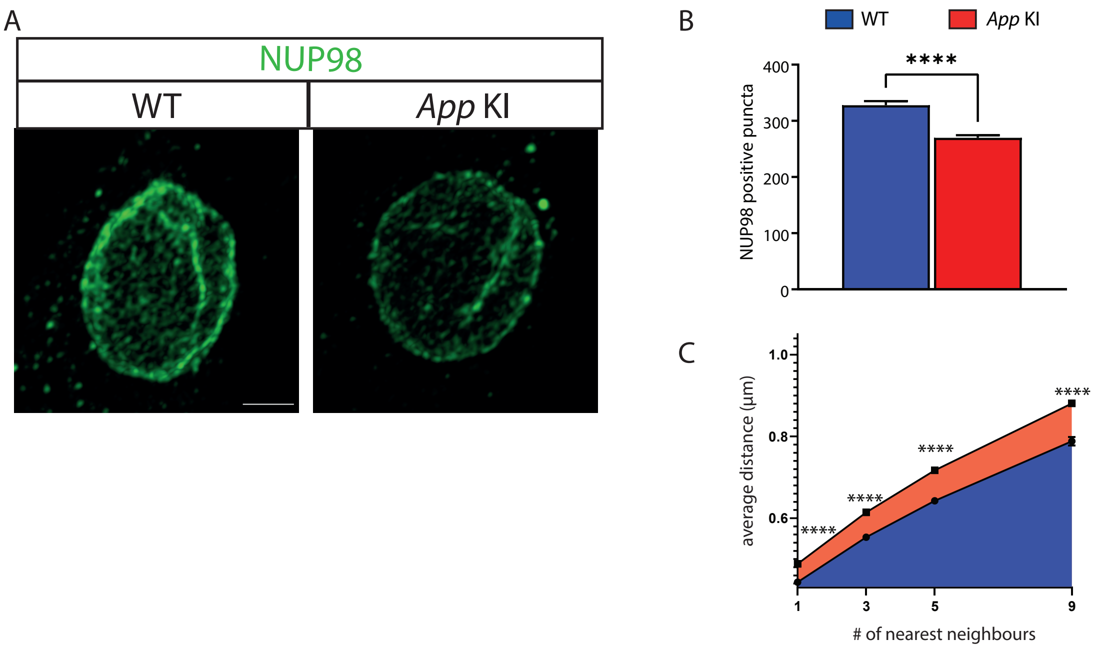


#### S7 Reduced NUP98 and NUP107 expression in APP/PS1 mice

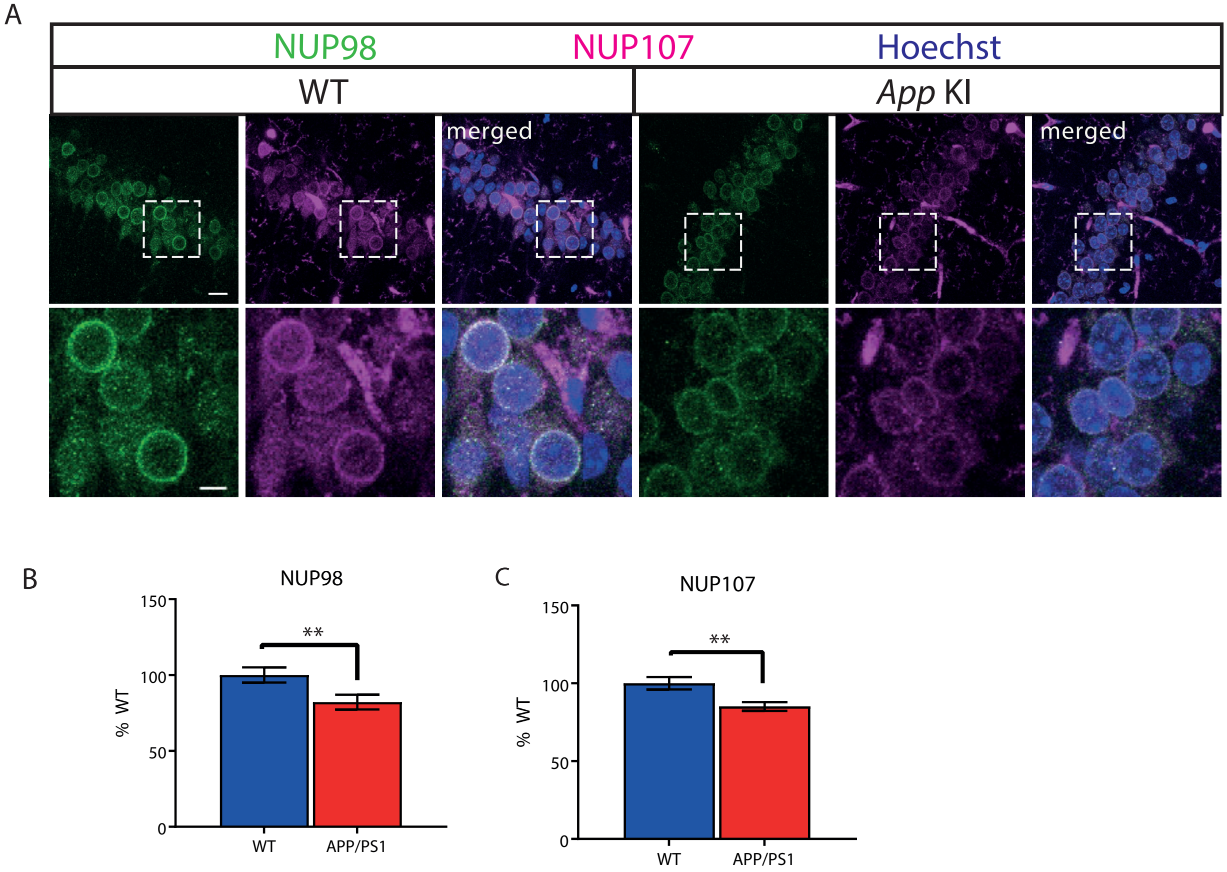

### SUPPLEMENTARY FIGURES

#### S8 Exogenous addition of Aβ oligomers induces loss of NPCs in WT neuronal nuclei

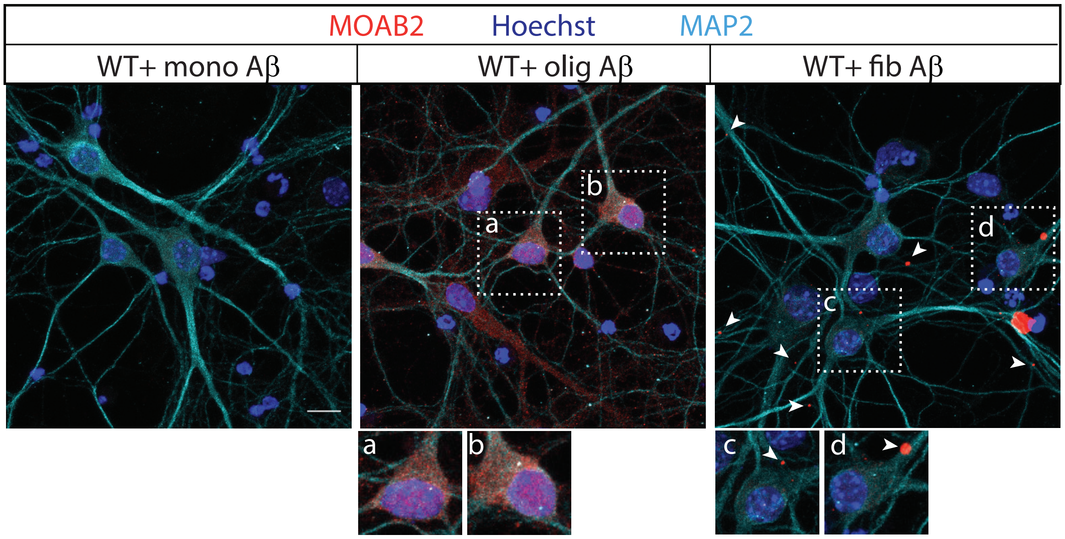


#### S9 4xEGFP accumulates in *App* KI neuronal nuclei

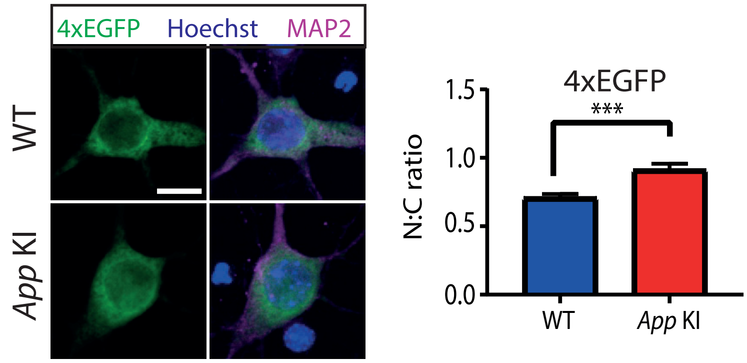


#### S10 NLS-4xEGFP accumulates in *App* KI neuronal cytoplasm

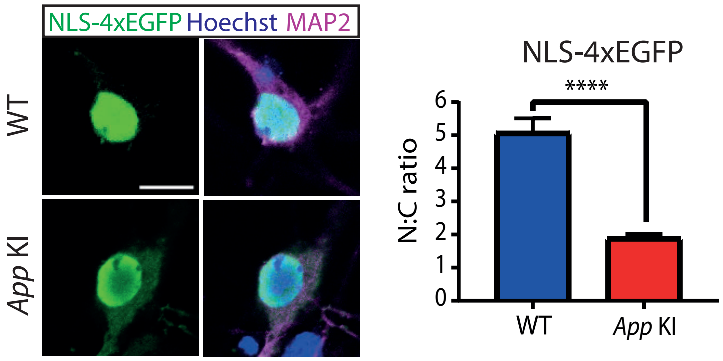


#### S11 LMB and GppNHP alters nuclear localization of CRTC1 and 4x-NLSGFP in cocultures

A

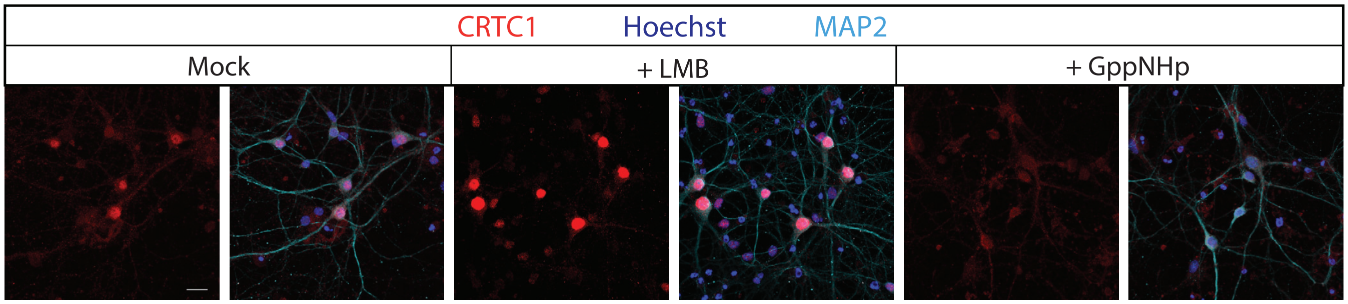


B

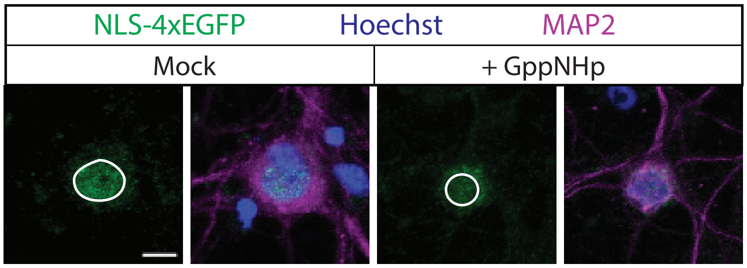

### SUPPLEMENTARY FIGURES

#### S12 Basal activation of necroptosis component proteins in *App* KI neurons enhances neuronal vulnerability

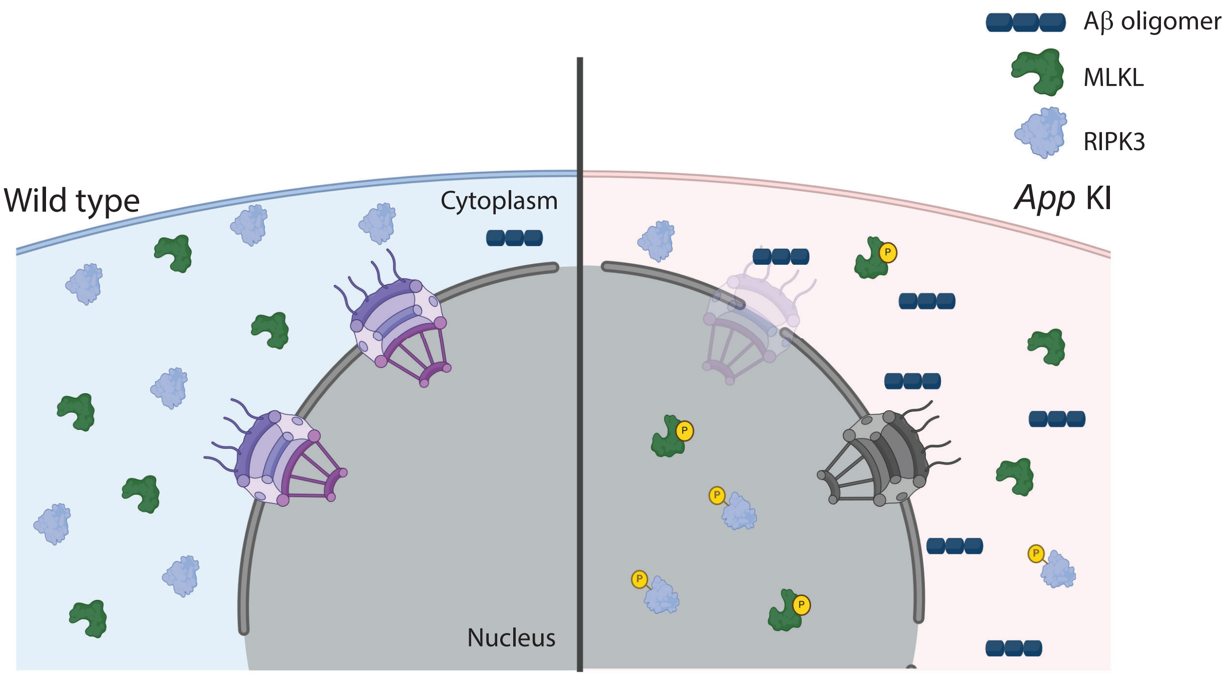


#### S13 Aβ in CA1 hippocampal pyramidal layer neurons in *App* KI (13-month old) mice

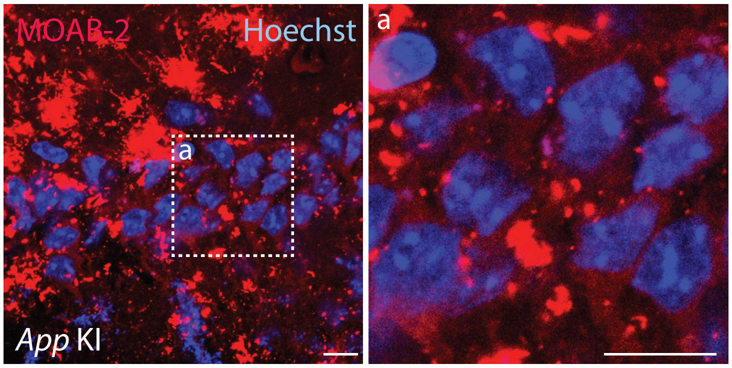


#### S14 Airyscan confocal micrograph of MOAB-2 antibody labeling of Aβ in neuronal soma

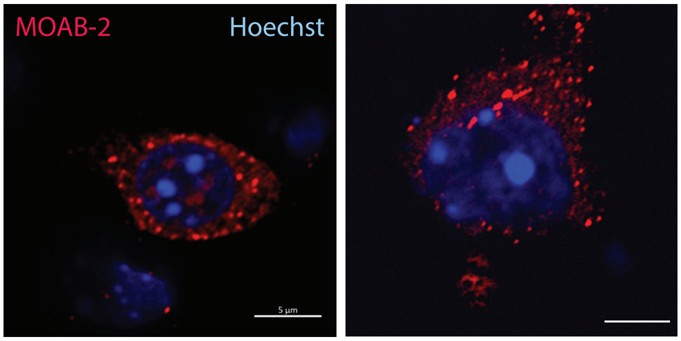


#### S15 Elevated nuclear CRTC1 in basal and TTX-silenced *App* KI neurons.

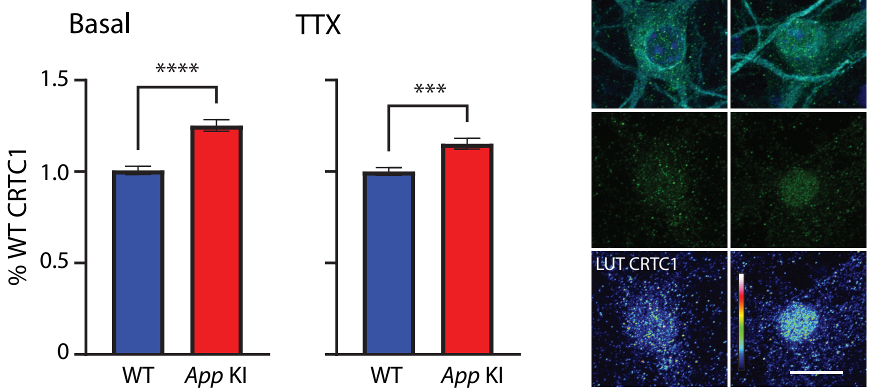
